# A *de novo* transcriptome assembly of the annelid worm *Hediste diversicolor*

**DOI:** 10.1101/2022.07.20.500767

**Authors:** Rodrigo Samico, André M. Machado, Marcos Domingues, Andreas Hagemann, Luísa Valente, Arne Malzahn, André Gomes-dos-Santos, Raquel Ruivo, Óscar Monroig, L. Filipe C. Castro

**Affiliations:** CIMAR/CIIMAR - Interdisciplinary Centre of Marine and Environmental Research, University of Porto, Avenida General Norton de Matos, S/N, 4450-208 Matosinhos, Portugal; FCUP - Department of Biology, Faculty of Sciences, University of Porto (U. Porto), Rua do Campo Alegre, Porto, Portugal; SINTEF Ocean, Environment and New Resources, Trondheim, Norway; ICBAS, Instituto de Ciências Biomédicas de Abel Salazar, Universidade do Porto, Porto, Portugal; Instituto de Acuicultura Torre de la Sal (IATS-CSIC), Castellón, Spain

**Keywords:** transcriptome, ecology, ragworm, Annelida

## Abstract

The development of Next-Generation Sequencing (NGS) technologies has revolutionized multiple fields of Biology. The ability to sequence DNA and RNA in an automated, parallel and low-cost approach, was key to boost the applicability and acquisition efficiency of *omics* resources. In this context, the availability of these tools has become indispensable towards our understanding of biodiversity, deepening our knowledge on the distinct complexity levels, from cells to ecosystems.

In this study we comprehensively characterised and annotated a whole-body transcriptome of *Hediste diversicolor*. This annelid worm species belongs to the family Nereididae and inhabits estuarine and lagoon areas on the Atlantic coasts of Europe and North America. Ecologically, this species plays an important role in benthic food webs. The ability of rework sediments through bioturbation activity makes this species essential to the estuarine mechanisms.

Here, we used Illumina next-generation sequencing technology, to sequence a total 105 million (M) paired-end (PE) raw reads and produce the first multi-tissue transcriptome assembly of an adult of *H. diversicolor*. This transcriptome contains 69,335 transcripts with a N50 transcript length of 2,313 bp and a BUSCO gene completeness of 97.7% and 96% (S: 88.2%; D: 7.8%) in Eukaryota and Metazoa lineage-specific profile libraries. Our findings offer a valuable resource for multiple biological studies using this species.

## 1. Introduction

The emergence of Next-Generation Sequencing (NGS) has provided the ability to sequence DNA at great speed and accuracy, allowing multiple approaches and innovative biological applications^1^. NGS debuted with the Human Genome Project^2^ and enabled the sequencing of DNA and RNA in a more automated and profitable way, leading to an exponential growth in our understanding and productivity^3^. Additionally, the consistent progression of transcriptomic datasets has provided innovative strategies to improve conservation and management of important species^4^.

The common ragworm *Hediste diversicolor* (O.F. Müller, 1776), occupies estuarine and lagoon areas of the Atlantic coasts of Europe and North America^5^. This aquatic annelid species belongs to the family Nereididae with great relevance due to its ecological role and commercial value^6^. These include the construction of burrows, through sediment reworking by bioturbation activity that intensify the sediment-water interface and create new micro-environments^7^, to their use as bait in sport and professional fishing or as food in aquaculture^8^. Given their ability to handle great fluctuations in salinity^9,10^ and temperature^11^, *H. diversicolor* specimens can be commonly found in a wide geographic range. Consequently, they are frequently used for toxicity assessment and biomonitoring of pollution levels^12,13^. However, this species is increasingly threatened due to its use as fishing bait and the lack of proper capture management^8^. This is further aggravated by the overlap between their reproductive peak and sport fishing season, during spring and summer; which, together with the fact they can only reproduce once in a lifetime,^14^ exacerbates the negative effects of the catch.

Here, using a combination of NGS approaches and a comprehensive bioinformatics workflow, we generated the first transcriptome for the common ragworm *H. diversicolor*. Despite the many relevant studies on this species, mainly focusing on its life-cycle, distribution and toxicology, no previous NGS dataset existed. With this data, we expect to provide novel and useful resources for future studies on this organism, to improve its conservation and sustainable exploitation.

## 2. Data description

### 2.1. Sampling, RNA extraction, and Illumina sequencing

An adult *H. diversicolor* individual was collected at Trondheim Fjord, in Leangbukta, Norway at 63.439151 N, 10.474605 E. The whole organism was stored in RNA later and frozen at −80°C until RNA extraction.

Total RNA extractions from the head and the middle body (reciprocal distance from the head and the tail) were performed with NZY Total RNA Isolation kit (NZYTech, Lda. - Genes and Enzymes) following the manufacturer’s recommendations. A DS-11 Series Spectrophotometer was used to evaluate RNA concentrations (ng/l) and quality (OD260/280 ratio values). Strand-specific libraries were built by Macrogen, Inc. These libraries included inserts between 250 and 300 bp, and were sequenced on the Illumina HiSeq4000 platform using 150 bp paired-end reads.

### 2.2. Processing of raw datasets and de novo assembly

Initially, we inspected the raw datasets (head and middle body) using the FastQC software (version 0.11.8) (http://www.bioinformatics.babraham.ac.uk/projects/fastqc/). Next, the Trimmomatic (version 0.38)^15^ software was used to quality-filter the raw reads and remove any Illumina adapters (Parameters: LEADING:5 TRAILING:5 SLIDINGWINDOW:4:5 MINLEN:36). In the end, the Rcorrector (version 1.0.3)^16^, a kmer-based error correction method, was executed with the default settings to correct any random sequencing errors. Importantly, this software corrected about 30 M of the 101 M reads in first sample and 28 M of the 108 M reads in the second one. Afterwards the FastQC was applied to verify the final quality of the dataset. Despite the conservatory approach used, few reads were discarded during the pre-processing stage and all final reads of both samples presented Phred scores higher than 30. After this process we deposited the raw reads in the NCBI database and can be consulted under the BioProject accession: PRJNA860539 (Table 1).

**Table 1.** MIxS information for transcriptome assembly for *Hediste diversicolor*.

|  | Value |
| --- | --- |
| Investigation_type | Eukaryote |
| Project_name | *Hediste diversicolor* transcriptome |
| Lat_lon | 63.439151 N, 10.474605 E |
| Geo_loc_name | Norway: Trondheim Fjord, Leangbukta |
| Collection_date | 10/07/2020 |
| Biome | Coastal sea water (ENVO:00002150) |
| Feature | Coastal water body (ENVO:02000049) |
| Material | Sea water (ENVO:00002150) |
| Env_package | RNAlater |
| Seq_meth | Illumina HiSeq |
| Assembly method | Trinity |
| Collector | Andreas Hagemann |
| Tissues | Head, Middle body |
| Maturity | Adult |
| Tissue sample size | 1 cm |
| Assembly | Trinity v.2.13.2 |
| BioProject | PRJNA860539 |

To build the *H. diversicolor* transcriptome assembly, we used the Trinity (version 2.13.2)^17^ software with the default settings and a single k-mer approach. Before run the Trinity software, both datasets were merged. Next, the merged file was submitted to Trinity, with the default parameters, and the initial transcriptome assembly produced (Assembly v0). To assess the v0 transcriptome assembly we used Transrate (version 1.0.1)^18^ software to assess general metrics of the assembly, the Benchmarking Universal Single-Copy Orthologs (BUSCO version 3.0.2)^19^ – with two lineage-specific profile libraries (Metazoa and Eukaryota) – to determine the gene completeness of the assembly and a read back mapping (RBM) approach to determine the percentage of clean reads present in the transcriptome assembly. To align the reads against the transcriptome assembly we used the Bowtie2 (version 2.3.5)^20^ software and to quantify the percentage of reads mapping we used the SAM stats (version 1.9)^21^. In this version of the transcriptome assembly, we found 280,456 transcripts, an n50 transcript length of 2,512 bp, an n90 transcript length of 361 bp, and a mean transcript length of 1081,46 bp (Table 2). The BUSCOs analysis presented a high level of gene completeness in both databases, 100% (S: 13.2%; D: 86.8%) and 99.8% (S: 14.3%; D: 85.5) in Eukaryota and Metazoa database, respectively (Table 2). The RBM analyses showed about 98.10% of the initial clean reads present in the transcriptome (Table 2).

**Table 2.** Statistics of the *Hediste diversicolor* transcriptome assembly versions, raw (V0), decontaminated (V1), final (V2).

| Raw Sequences | *Hediste diversicolor* |  |  |
| --- | --- | --- | --- |
| Reads used in assembly | 105,011,718 |  |  |
| Quality Metrics of Assembly | Assembly v0 | Assembly v1 | Assembly v2 |
| Number of transcripts | 280,456 | 272,156 | 69,335 |
| N50 transcript length (bp) | 2,512 | 2,487 | 2,313 |
| N90 transcript length (bp) | 361 | 358 | 556 |
| Transcriptome length (mb) | 330,6 | 298,9 | 96,8 |
| Mean transcript length (bp) | 1081,46 | 1071,22 | 1335,81 |
| Longest transcript | 22,036 | 22,036 | 22,036 |
| Number of transcripts over 1K nt | 76,694 | 73,343 | 25,074 |
| GC % | 0.40 | 0.40 | 0.39 |
| Rate Mapping % | 98.10 | 96.47 | 91.69 |
| **BUSCO Complete (euk/met) %** | 100 | 99.6 | 97.7 |
| Single | 13.2/14.3 | 19.1/17.4 | 89.8/88.2 |
| Duplicated | 86.8/85.5 | 80.5/81.7 | 7.9/7.8 |
| Fragmented | 0.0/0.1 | 0.3/0.5 | 2.0/2.7 |
| Missing | 0.0/0.1 | 0.1/0.4 | 0.3/1.3 |
| **Annotation** | - | - | - |
| Number of transcripts with ORF | - | - | 32,006 |
| Transcripts with Blast hit in at least one database | - | - | 25,453 |
| Transcripts with KEGG Pathways | - | - | 1,168 |
| Transcripts with Interpro hit | - | - | 21,745 |
| Transcripts with PFAM | - | - | 19,088 |
| Transcripts with GO terms | - | - | 13,402 |

### 2.3. Post-assembly processing stage

After the transcriptome assembly and initial validations, we ran the decontamination analysis. In summary, we used the nucleotide database of NCBI (nt-NCBI; downloaded on 17/02/2022) and UniVec (downloaded on 17/02/2022) database to identify and remove possible sources of contamination (e.g., vectors, adapters, biological contamination). These analyses were done using blast-n (version 2.12.0)^22^ versus the nt-NCBI dataset with the settings (-num_threads 15 -max_target_seqs 1 - max_hsps 1) and versus the Univec dataset with the parameters (-reward 1 -penalty -5 -gapopen 3 -gapextend 3 -dust yes -soft_masking true -evalue 700 -searchsp 1750000000000 -outfmt 6 -num_threads 15 -perc_identity 90; and minimum alignment length of 100 bp). To the nt-NCBI searches, all the transcripts with the best match hits out of the Lophotrochozoa taxon (Taxonomy ID: 1206795) were considered contaminations and were excluded from the v0 transcriptome assembly. The remaining contigs were preserved in the transcriptome and used in subsequent analyses, whether they had or had not match hits in the Lophotrochozoa taxon. All transcripts having a match hit in the Univec database were deemed exogenous to the *H. diversicolor* transcriptome and deleted from the dataset. In the end of this process, we obtained the decontaminated assembly (Assembly v1) (Table 2). To remove redundancy of the datasets we executed the Corset tool^23^. The program removes transcripts with less than ten read counts (spurious transcripts) and hierarchically cluster the remaining transcripts based on the fraction of common reads multi-mapped to the transcriptome. Next, the FetchClust (version 1) (https://github.com/Adamtaranto/Corset-tools/blob/master/fetchClusterSeqs.py) with the parameter (–longest) was used to select the longest transcripts from each cluster (Assembly v2).

To perform the assessment of v1 and v2 versions of the transcriptome assembly we used the same strategy used to v0. While the TransRate analyses showed an n50 transcript length of 2,487 (v1) and 2,313 bp (v2) and n90 transcript length of 358 (v1) and 556 bp (v2) the BUSCOs analysis showed 99.6% (v1) and 97.7% (v2) of gene completeness in Eukaryota database and 99.1 (v1) and 96% (v2) against Metazoa database. Despite the large reducing in the number of transcripts from v1 (272,156) to v2 (69,335), the BUSCO analyses showed that almost no gene content was lost. The redundance removal step affected mainly transcripts with small length (0-800bp) and consequently the mean length of the transcriptome, 1071,22 (v1) and 1335,81 bp (v2) (Figure 1a). Additionally, the number of isoforms also was slightly affected Figure 2b.

**Figure 1.**
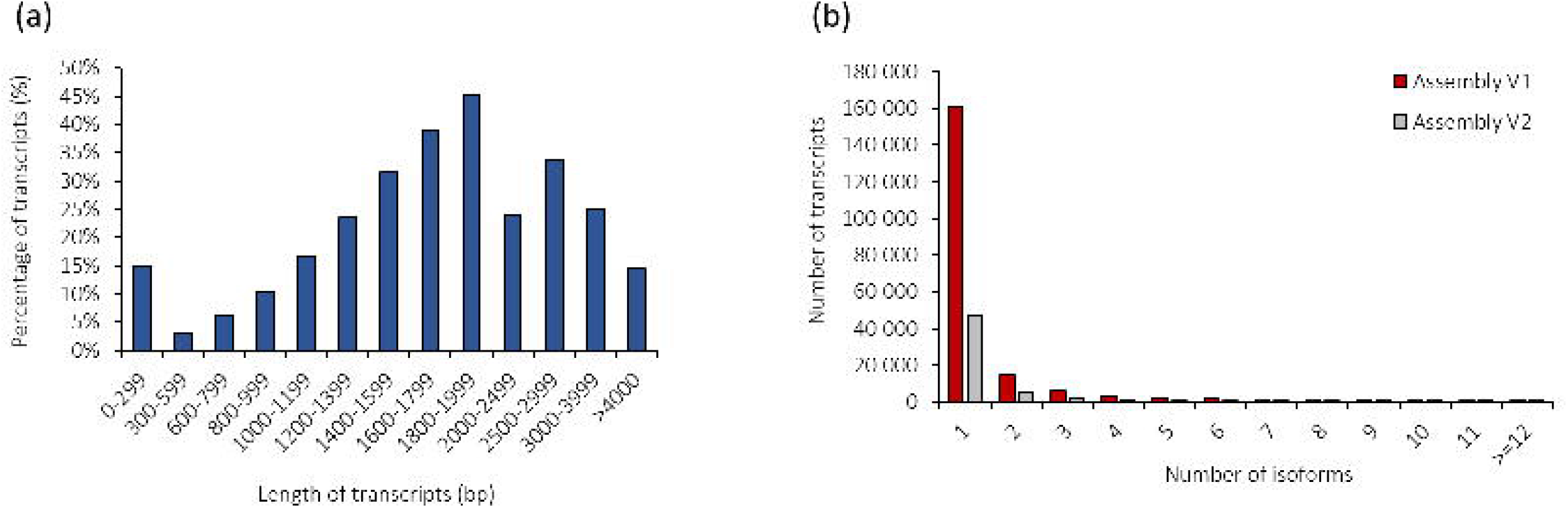
Quality assessment of the transcriptome of *Hediste diversicolor*. a) Transcript length distribution b) Number of isoforms per cluster of transcripts.

**Figure 2.**
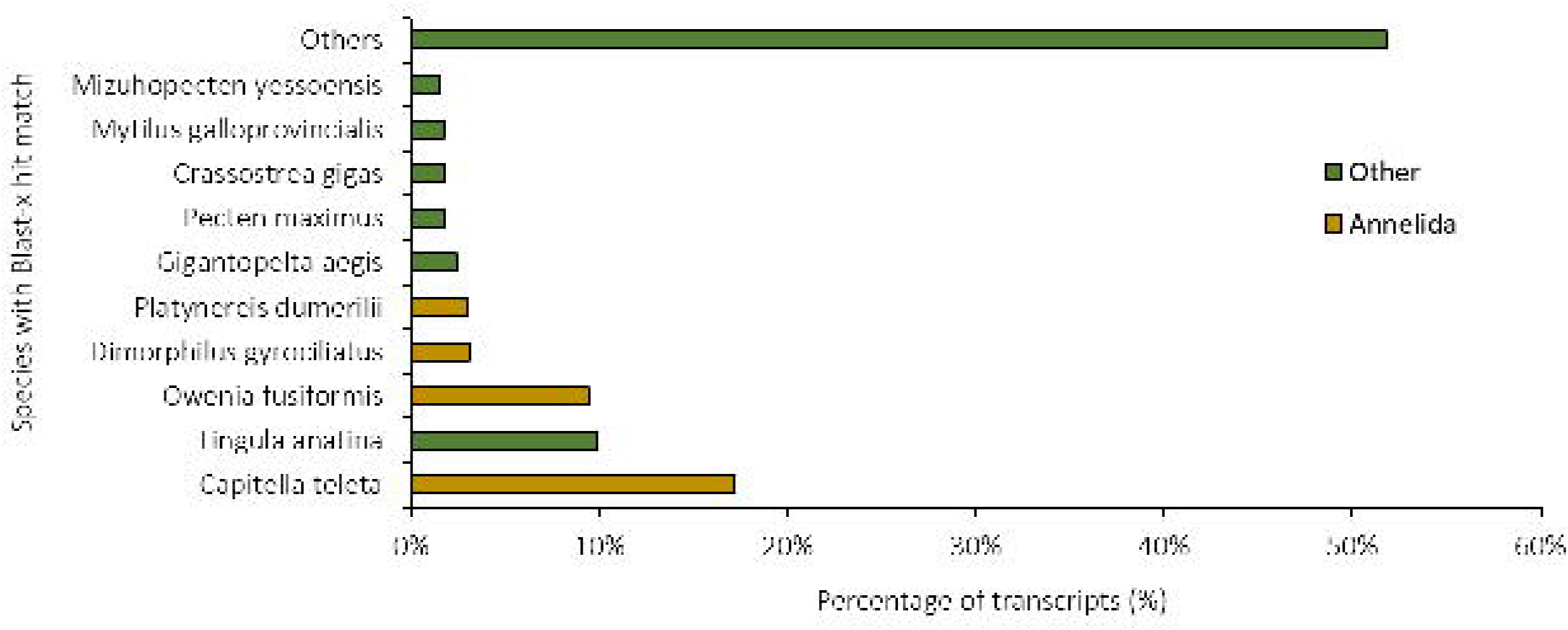
Blast-x analysis of the transcriptome assembly of *Hediste diversicolor*.

### 2.4. Transcriptome annotation

The annotation of the *H. diversicolor* transcriptome was subdivided in two steps, structural and functional annotation. First, to perform the structural annotation we executed the TransDecoder (version 5.3.0) (https://github.com/TransDecoder/TransDecoder/). Transdecoder allowed us to determine possible coding regions within transcript sequences and produce the open reading frames (ORFs). To do that, we used both homology and protein domain searches - homologies searches with Blast-p^24^ (version 2.12.0) tool in UniProtKB/SwissProt database^25^ and domain searches with hmmscan software of hmmer package (version 3.3.2) in PFAM database^26^.

To perform the functional annotation, we used a set of tools. Initially Gtf/Gff Analysis Toolkit (AGAT) (version 0.8.0)^27^ allowed us to uniformize and produce the initial structural annotation file (in gff3 format) using the transcriptome assembly file (.fasta) and the Trandecoder output file (.gff). In the end, the protein and transcript fasta files were obtained with names properly uniformized and formatted. In the last phase of the annotation, the functional annotation, we applied the InterProScan tool (version 5.44.80)^28^ and Blast-n/p/x searches in several databases. While the InterPro and NCBI protein databases were queried for the proteins per species using the Blast-p/x tool of DIAMOND software (version 0.9.24)^29^, the NCBI-nt and NCBI-nr databases were searched for transcripts using the Blast-n/x tool of DIAMOND software. In the end, the AGAT program combined all blast (outfmt6 files) and InterProScan (tsv file) outputs into the gff3 annotation file. The top 10 species with the best blast-x matching against *H. diversicolor* transcriptome can be consulted in Figure 2. This distribution clearly reflects the lack of genomic resources in Annelida phylum. Moreover, in these 10 species only three species, *Capitella teleta, Owenia fusiformes* and Dimorphilus gyrociliatus, equivalent to 29,75 % of the total hits, belong to Annelida.

## 3. Conclusions

We generated the first transcriptome assembly for the common ragworm *Hediste diversicolor*. The results and resources present here provide a significant advancement in the study of this group of organisms.

## Acknowledgments

This research was partially supported by the ERA-NET BlueBio COFUND Project SIDESTREAM [Grant ID 68], co-funded through national funds provided by FCT [BLUEBIO/0005/2019]; Agencia Estatal de Investigación [PCI2020-111960], NRC – Norwegian Research Council [#311701]. A.G.S. was funded by the Portuguese Foundation for Science and Technology (FCT) under the Grant [SFRH/BD/137935/2018]. Additional strategic funding was provided by FCT [UIDP/04423/2020].

## Notes

### Competing Interest Statement

The authors have declared no competing interest.

## References

1. Zhang, J., Chiodini, R., Badr, A. & Zhang, G. The impact of next-generation sequencing on genomics. J. Genet. Genomics Yi Chuan Xue Bao 38, 95–109 (2011).

2. Gibbs, R. A. The Human Genome Project changed everything. Nat. Rev. Genet. 21, 575–576 (2020).

3. Behjati, S. & Tarpey, P. S. What is next generation sequencing? Arch. Dis. Child. Educ. Pract. Ed. 98, 236–238 (2013).

4. Connon, R. E., Jeffries, K. M., Komoroske, L. M., Todgham, A. E. & Fangue, N. A. The utility of transcriptomics in fish conservation. J. Exp. Biol. 221, jeb148833 (2018).

5. Virgilio, M. & Abbiati, M. Habitat discontinuity and genetic structure in populations of the estuarine species Hediste diversicolor (Polychaeta: Nereididae). Estuar. Coast. Shelf Sci. 61, 361–367 (2004).

6. Einfeldt, A. L., Doucet, J. R. & Addison, J. A. Phylogeography and cryptic introduction of the ragworm *Hediste diversicolor* (Annelida, Nereididae) in the Northwest Atlantic. Invertebr. Biol. 133, 232–241 (2014).

7. Gillet, P., Mouloud, M. & Mouneyrac, C. The key role of the species Hediste Diversicolor (polychaeta, nereididae) in estuarine ecosystems. in (2018).

8. Scaps, P. A review of the biology, ecology and potential use of the common ragworm Hediste diversicolor (O.F. Müller) (Annelida: Polychaeta). Hydrobiologia 470, 203–218 (2002).

9. Wolff, W. J. The estuary as a habitat an analysis of data on the soft-bottom Macrofauna of the Estuarine area of the rivers rhine, Meuse, and Scheldt. Zool. Verh. 126, 1–242 (1973).

10. Neuhoff, H.-G. Influence of Temperature and Salinity on Food Conversion and Growth of Different Nereis Species (Polychaeta, Annelida). Mar. Ecol. Prog. Ser. 1, 255–262 (1979).

11. Newell, R. C. & Branch, G. The Influence of Temperature on the Maintenance of Metabolic Energy Balance in Marine Invertebrates. in Advances in Marine Biology vol. 17 329–396 (1980).

12. Ozoh, P. T. E. The importance of adult Hediste (Nereis) diversicolor in managing heavy metal pollution in shores and estuaries. Environ. Monit. Assess. 21, 165–171 (1992).

13. Buikema, A. L., Buikema, J. & Cairns, J. Aquatic Invertebrate Bioassays. (ASTM International, 1980).

14. Olive, P. J. W. & Garwood, P. R. G. Gametogenic cycle and population structure of Nereis (Hediste) diversicolor and Nereis (Nereis) pelagica from north-east England. J. Mar. Biol. Assoc. U. K. 61, 193–213 (1981).

15. Bolger, A. M., Lohse, M. & Usadel, B. Trimmomatic: a flexible trimmer for Illumina sequence data. Bioinformatics 30, 2114–2120 (2014).

16. Song, L. & Florea, L. Rcorrector: efficient and accurate error correction for Illumina RNA-seq reads. GigaScience 4, s13742-015-0089-y (2015).

17. Haas, B. J. et al. De novo transcript sequence reconstruction from RNA-seq using the Trinity platform for reference generation and analysis. Nat. Protoc. 8, 1494–1512 (2013).

18. Smith-Unna, R., Boursnell, C., Patro, R., Hibberd, J. M. & Kelly, S. TransRate: reference-free quality assessment of de novo transcriptome assemblies. Genome Res. 26, 1134–1144 (2016).

19. Simão, F. A., Waterhouse, R. M., Ioannidis, P., Kriventseva, E. V. & Zdobnov, E. M. BUSCO: assessing genome assembly and annotation completeness with single-copy orthologs. Bioinformatics 31, 3210–3212 (2015).

20. Langmead, B. & Salzberg, S. L. Fast gapped-read alignment with Bowtie 2. Nat. Methods 9, 357–359 (2012).

21. Lassmann, T., Hayashizaki, Y. & Daub, C. O. SAMStat: monitoring biases in next generation sequencing data. Bioinformatics 27, 130–131 (2011).

22. Altschul, S. F. et al. Gapped BLAST and PSI-BLAST: a new generation of protein database search programs. Nucleic Acids Res. 25, 3389–3402 (1997).

23. Davidson, N. M. & Oshlack, A. Corset: enabling differential gene expression analysis for de novoassembled transcriptomes. Genome Biol. 15, 410 (2014).

24. Altschul, S. F., Gish, W., Miller, W., Myers, E. W. & Lipman, D. J. Basic local alignment search tool. J. Mol. Biol. 215, 403–410 (1990).

25. Apweiler, R. et al. UniProt: the Universal Protein knowledgebase. Nucleic Acids Res. 32, D115–D119 (2004).

26. Mistry, J. et al. Pfam: The protein families database in 2021. Nucleic Acids Res. 49, D412–D419 (2021).

27. Dainat J., Hereñú D., Pucholt P. AGAT: Another Gff Analysis Toolkit to handle annotations in any GTF/GFF format. Zenodo.

28. Blum, M. et al. The InterPro protein families and domains database: 20 years on. Nucleic Acids Res. 49, D344–D354 (2021).

29. Buchfink, B., Xie, C. & Huson, D. H. Fast and sensitive protein alignment using DIAMOND. Nat. Methods 12, 59–60 (2015).

